# Peaceful Queen Succession in the Naked Mole Rat

**DOI:** 10.64898/2026.01.09.698689

**Authors:** Shanes C. Abeywardena, Alexandria M. Schraibman, Victor Delgado Cuevas, Janelle S. Ayres

**Affiliations:** Animal Resources Department, Salk Institute for Biological Studies, 10010 N. Torrey Pines Road, La Jolla, CA 92037, USA; Molecular and Systems Physiology Lab, Salk Institute for Biological Studies, 10010 N. Torrey Pines Road, La Jolla, CA 92037, USA; Howard Hughes Medical Institute, Salk Institute for Biological Studies, 10010 N. Torrey Pines Road, La Jolla, CA 92037, USA

## Abstract

The eusocial naked mole rat exhibits an extreme reproductive skew involving a single queen that monopolizes breeding through behavioral dominance. When reproductive suppression of subordinates is lifted due to removal or death of the queen, subordinate females compete to assume the reproductive role, resulting in intense aggression and intracolony wars. This aggressive reproductive strategy is thought to confer several advantages under stable environmental conditions. However, reliance on a single reproductive female has the potential to create vulnerabilities when challenged with stressors or unstable environments. Thus, in some contexts, non-violent reproductive flexibility may be beneficial. Here, we report a longitudinal study of a captive naked mole rat colony identifying an alternative, mechanistically distinct mode of queen succession in response to external stressors that impair queen reproduction without social disturbance. Elevated colony density was sufficient to impair pup survival but did not alleviate reproductive suppression or trigger aggression in the colony. By contrast, relocating the colony to a new facility caused a long-term pause in the queen’s reproduction, leading to the sequential emergence of her daughters as additional breeders in the absence of fighting or aggression. This resulted in a transient period of peaceful plural breeding with one daughter and ultimately resulted in another daughter assuming the primary reproductive role for the colony. Together our findings support a model in which reproductive ascension is permissive and socially tolerated when the reigning queen’s reproductive function is impaired, thereby expanding the mechanistic framework of naked mole rat eusociality to include peaceful, fertility-based pathways of queen succession.

## Introduction

Across the animal kingdom, numerous mating systems have evolved, ranging from monogamy and polygyny, to cooperative breeding and eusociality. In most mammals, reproduction is typically distributed in an unequal manner with polygyny, where one male mates with multiple females, being the most common mating system (*1*). At the extreme end of the mating spectrum is eusociality, which is characterized by a division of labor into reproductive animals and non-reproductive workers that support colony function, maintenance and defense (*2*). The best appreciated examples of eusociality involve social insects including ants, bees and termites (*3*). Among mammals, the naked mole rat, *Hetercephalus glaber*, also exhibits eusociality (*4*). Naked mole rat colonies are typically organized around a single breeding female called the queen, with one to three breeding males, while the majority of colony members remain non-reproductive (*5*). Reproduction in this system is closely tied to social dominance, with the queen actively suppressing ovulation and reproductive attempts by subordinate females through behavioral, pheromonal and possibly physiological mechanisms (*6*). If the queen dies or is removed from the colony, the reproductive suppression imposed on subordinate females is lifted, and subordinate females compete to assume the reproductive role, resulting in intense aggression and intracolony wars (*5, 7*). In addition to this classic succession pathway, new queens can arise through other mechanisms. Within the natal colony, a subordinate female may attempt a coup, challenging the reigning queen (*5*). If successful, she will assume reproductive dominance (*5*). Subordinates may also engage in dispersal behavior, leaving the natal colony to seek a mate and establish a new colony elsewhere, becoming queen of the new group (*8*). Finally in captive settings, a subordinate female can be removed from her natal colony and paired with a male in a separate enclosure, where she can mate and become a reproductive queen (*7*).

At the colony level, the rigid aggressive strategy for naked mole rat reproduction has been proposed to confer several fitness advantages under stable conditions (*5*). However, there are several potential evolutionary disadvantages to this strategy. Reliance on a single reproductive female limits a colony’s ability to hedge against unpredictable events such as disease, predation or environmental disruptions where plural breeding could buffer against a failure of any single reproductive female (*9, 10*). Furthermore, the reliance on intense aggression and physical dominance to enforce reproductive suppression can carry additional costs including the risk of injury and substantial energetic costs (*11*). High levels of within-colony conflict may also undermine social cohesion. Moreover, strict suppression maintains reproductively competent subordinate females in a non-breeding state even when environmental opportunities for dispersal or cooperative plural breeding might enhance long term fitness (*8*). Thus, although strict eusociality provides stability under predictable and stable conditions, it introduces significant tradeoffs. This suggests that some level of reproductive plasticity that allows for flexible modulation of social structure in response to internal or external cues, may have evolved as a contingent alternative.

Here we describe our longitudinal study of a laboratory naked mole rat colony in which there was peaceful queen succession. Using environmental stressors known to impair reproduction in other rodents, we disrupted the reproductive success of an established queen without triggering aggression or social instability. Increased colony density did not prevent the queen from conceiving or giving birth but markedly reduced postnatal litter survival without lifting the reproductive suppression in subordinates. Relocating the colony to a new vivarium produced more severe impairment, temporarily halting the queen’s reproduction. Under these conditions, we revealed non-violent, asynchronous but partially overlapping pregnancies with the queen and a subordinate female. Upon removal of this subordinate female, a second subordinate subsequently became the sole reproductive female for the colony, again in the absence of aggression or fighting. Together our findings demonstrate that naked mole rat colonies are capable of non-aggressive, cooperative shifts in reproductive roles, revealing flexibility in naked mole rat reproductive biology.

## Results

### Establishment of a reproductive baseline for the Amigos colony

An overview of the study and data summary are shown in **Figure 1**, **Table 1** and **Supplemental Figure 1**. The Amigos colony was established in our facility on July 17, 2019. The family consisted of six animals: one reproductive queen, Teré (unknown age), a single male named Paquíto (unknown age), along with their first litter, born on May 8, 2019. This litter was comprised of five pups, but only four (three females and one male) survived. Between September 12-19, 2019, Teré exhibited a 4-gram weight gain and displayed physical features consistent with pregnancy (**Figure 2A** and **Supplemental Figure 1A**). We confirmed the presence of fetuses by ultrasound (**Figure 2B**). On September 26, 2019, she gave birth to her second litter, consisting of seven pups, all of which survived, increasing the colony size to 13 animals (**Figure 1**, **Table 1**, **Figure 2C-D** and **Supplemental Figure 1B-C**). During the first year following colony establishment (July 17, 2019-August 5, 2020), Teré produced an additional four litters at regular intervals of 76-81 days (**Figure 1**, **Table 1**, **Figure 2C, E** and **Supplemental Figure 1B, D**). Litter sizes ranged from 6-10 pups with 100% survival, with the exception of her fourth litter, which yielded a single pup that did not survive (**Figure 1**, **Table 1**, **Figure 2C** and **Supplemental Figure 1B**). In each case, predictable prepartum weight gain facilitated our identification of pregnancy status of the queen (**Figure 2A** and **Supplemental Figure 1A**). Across this baseline period, we observed no aggression or fighting within the colony, and no signs of injuries consistent with fighting (**Table 2**). Taken together, these observations establish that under stable housing conditions and social structure, Queen Teré demonstrated regular, predictable and successful reproductive output, consistent with a healthy dominant breeding female in the colony. Because survival to adulthood requires appropriate maternal and colony level care, these early results also confirm that successful reproduction included both parturition and postnatal survival supported by the colony.

**Figure 1.**
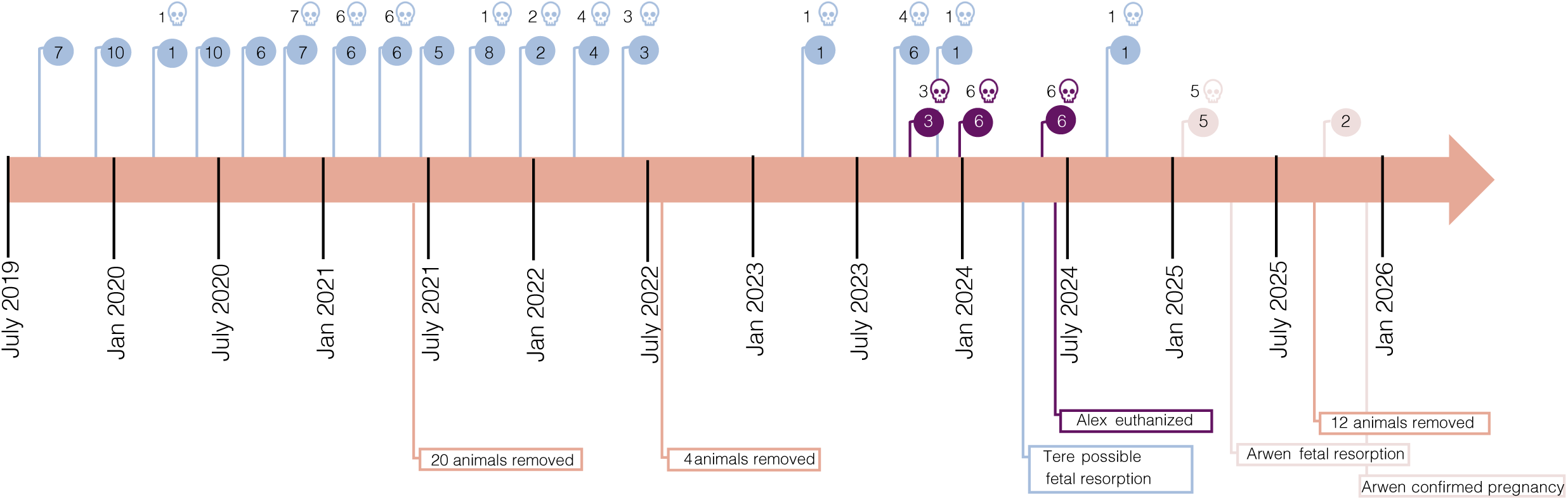
Summary of events in the Amigo Colony during the study time period. Circles indicate litters with the number of pups born in each litter noted inside the circle. Skull icons listed above the circles indicate the number of pups that died prior to one month of age. Blue circles indicate Queen Teré. Purple circles indicate Alexandria and peach circles indicate Arwen. Other important events including removal of animals, fetal reabsorption events and Alexandria’s death are noted.

**Figure 2.**
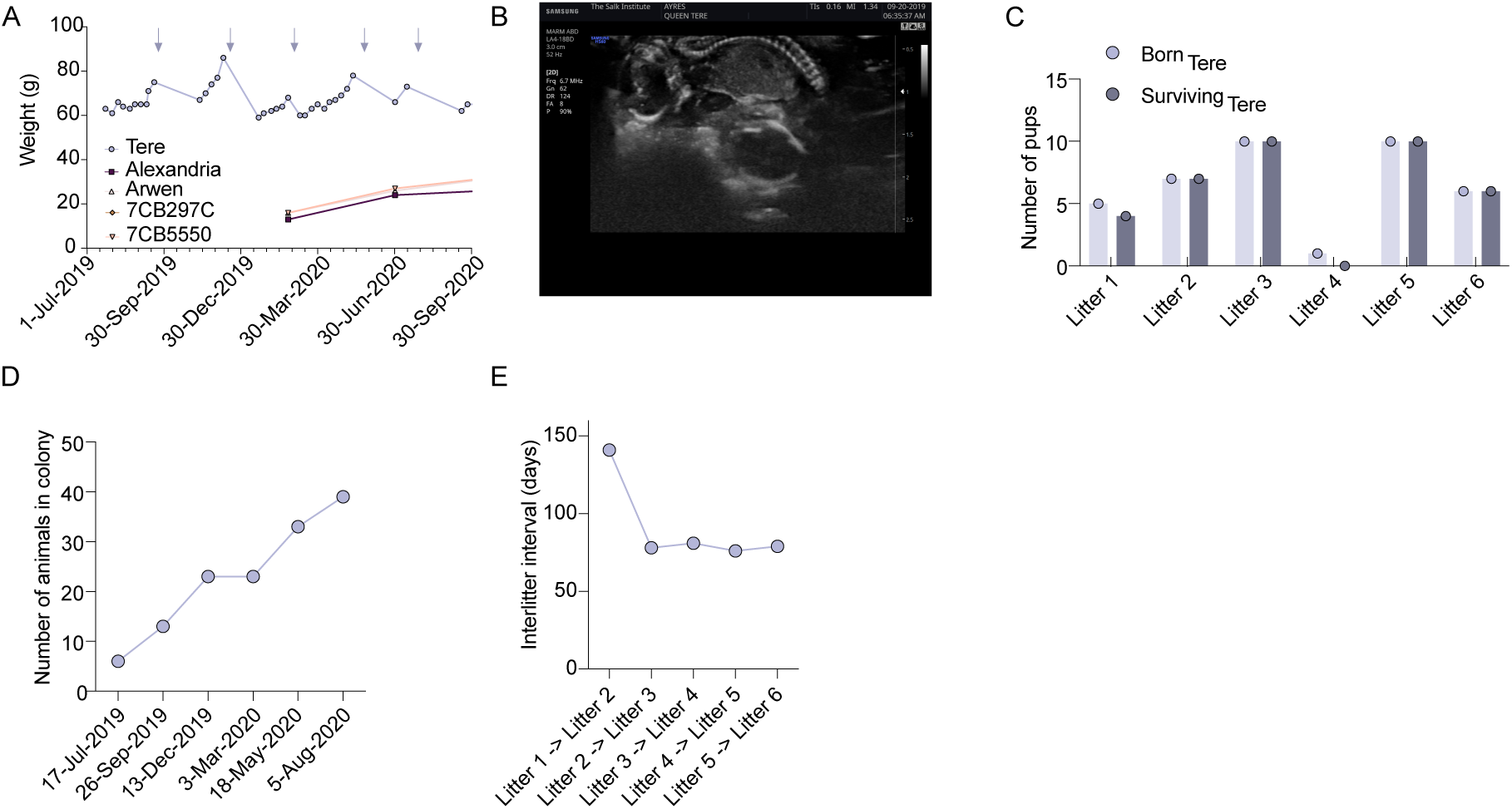
Establishment of a Reproductive Baseline for the Amigos Colony. (**A**) Body weight of Queen Teré and four of her daughters born in Queen Teré’s December 13, 2019 litter. Blue arrows indicate Queen Teré’s litters. (**B**) Ultrasound on Queen Teré performed on September 20, 2019. (**C**) The number of pups born and the number of surviving pups past one month of age for the indicated litters. (**D**) The number of animals in each colony one month following the noted litter dates. (**E**) Interlitter interval days between the indicated litters for the colony. Data shown in panels **A**, **C**, **D** and **E** also shown in **Supplemental** Figure 1A.

**Table 1.** Summary of reproductive data.

| Pregnancy Number* | Female | Litter Birthdate | Litter Size | Number of Survivors | Interlitter Interval (days)<sup>5</sup> | Total number of animals in colony<sup>6</sup> |
| --- | --- | --- | --- | --- | --- | --- |
| 1<sup>#</sup> | Tere | 5/8/2019 | 5 | 4 | / | 6 |
| 2 | Tere | 9/26/2019 | 7 | 7 | 141 | 13 |
| 3 | Tere | 12/13/2019 | 10 | 10 | 78 | 23 |
| 4 | Tere | 3/3/2020 | 1 | 0 | 81 | 23 |
| 5 | Tere | 5/18/2020 | 10 | 10 | 76 | 33 |
| 6 | Tere | 8/5/2020 | 6 | 6 | 79 | 39 |
| 7 | Tere | 10/25/2020 | 7 | 0 | 81 | 39 |
| 8 | Tere | 1/13/2021 | 6 | 0 | 80 | 39 |
| 9 | Tere | 4/4/2021 | 6 | 0 | 81 | 39 |
| 10 | Tere | 6/24/2021 | 5 | 5 | 81 | 24<sup>1</sup> |
| 11 | Tere | 9/13/2021 | 8 | 7 | 81 | 31 |
| 12 | Tere | 12/3/2021 | 2 | 0 | 81 | 31 |
| 13 | Tere | 2/21/2022 | 4 | 0 | 80 | 31 |
| 14 | Tere | 5/12/2022 | 3 | 0 | 80 | 31 |
| 15 | Tere | 4/9/2023 | 1 | 0 | 332 | 27<sup>2</sup> |
| 16 | Tere | 8/28/2023 | 6 | 2 | 141 | 29 |
| 17 | Alexandria (suspected) | 10/10/2023 | 3 | 0 | 43 | 29 |
| 18 | Tere | 11/16/2023 | 1 | 0 | 37 | 29 |
| 19 | Alexandria | 12/30/2023 | 6 | 0 | 44 | 29 |
| 20 | Tere | 4/23/2024 ultrasound possible fetal resorption | / | / | / | 29 |
| 21 | Alexandria | 5/23/2024 | 6 | 0 | 145 | 29 |
| 22 | Tere | 9/9/2024 | 1 | 0 | 109 | 28<sup>3</sup> |
| 23 | Arwen | 1/11/2025 | 5 | 0 | 124 | 28 |
| 24 | Arwen | 4/11/2025 ultrasound fetal resorption | / | / | / | 28 |
| 25 | Arwen | 10/5/2025 | 2 | 2 | 267 | 17<sup>4</sup> |
| 26 | Arwen | 12/31/25 ultrasound confirmed pregnancy |  |  |  |  |

\*Pregnancy confirmed by ultrasound and/or birth
#born at CUNY
<sup>1</sup> 20 animals removed on 6/21/21 to reduce colony density. See Supplemental Table 1
<sup>2</sup> 4 animals removed from colony between 7/8/22-8/18/22. See Supplemental Table 1
<sup>3</sup> Alexandria euthanized - uterine torsion

**Table 2.** Summary of injuries and aggressive behavior.

| Animal ID | Birthdate | Weight (g) | Sex | Date | Observations | Outcome | Aggressive behavior observed? |
| --- | --- | --- | --- | --- | --- | --- | --- |
| 7AA8C55 | 7/18/2019 | 39 | M | 4/20/2020 | Scabbing on muzzle | Resolved 4/21/2020 | No |
| 7BF3A7D | 9/26/2019 | 30 | F | 12/17/2022 | Facial bruising | Resolved 12/20/2022 | No |
| Tere | Unknown | 64 | F | 1/28/2023 | 2mm scab perineal region | Resolved 2/1/2023 | No |
| 7CB297C | 12/13/2019 | 45 | F | 2/6/2025 | Suspected bite wound and blood on face | Resolved wound on 2/7/2025 | No |
| 7BDCE23 | 7/18/19 | 53 | F | 6/30/25 | Animal found dead | Examination revealed no injuries or wounds. Cecal enlargement - possible postmortem effect | No |

### Elevated colony density impairs postnatal litter viability

To test our hypothesis that naked mole rat queen succession can also proceed peacefully under certain conditions, we first sought to identify perturbations capable of interrupting the queen’s reproductive output without her removal (intentional removal or death) and that do not trigger aggression, thereby allowing us to examine how the colony responds when reproductive failure occurs in the absence of social collapse. Colony density is known to impair reproductive performance in many rodent species by reducing fertility, pup survival, or both (*12–15*). Importantly, in naked mole rats, high density conditions have not been shown to provoke dominance challenges or social instability. We therefore tested the hypothesis that increased colony density would disrupt queen reproductive output without inducing colony aggression.

After Teré’s sixth litter (August 5, 2020), the Amigos colony reached 39 animals. Teré subsequently produced three additional litters of 6-7 pups at typical interlitter intervals of 80-81 days and with the predictable prepartum weight gain (**Figure 1**, **Table 1**, **Figure 3A-C** and **Supplemental Figure 1A-B, D**). Although she continued to become pregnant and deliver litters of normal size, 100% of her pups in each of these litters died shortly after birth, indicating a marked reduction in postnatal survival despite preserved fertility (**Figure 1**, **Table 1**, **Figure 3B** and **Supplemental Figure 1B**). We did not observe any fighting, aggressive behavior or injuries consistent with fighting in the colony during this time period (**Table 2**). To test whether reducing the colony density would restore pup vitality, we removed 20 animals during Teré’s pregnancy with her tenth litter (on June 21, 2021) (**Supplemental Table 1**), reducing the colony density to 19 animals (**Figure 1**, **Table 1**, **Figure 3D** and **Supplemental Figure 1C**). Following this intervention reproductive outcomes temporarily improved, with Teré producing two successive litters with the majority of pups surviving (100% and 87.5% respectively) (**Figure 1**, **Table 1**, **Figure 3B** and **Supplemental Figure 1B**). After colony density increased to 31 animals, Teré subsequently produced three consecutive litters at normal intervals but with reduced litter size (2-4 pups) and 100% mortality (**Figure 1**, **Table 1**, **Figure 3B-C** and **Supplemental Figure 1B-D**). As before, we did not observe aggression, fighting or wounds consistent with fighting during this time period (**Table 2**).

**Figure 3.**
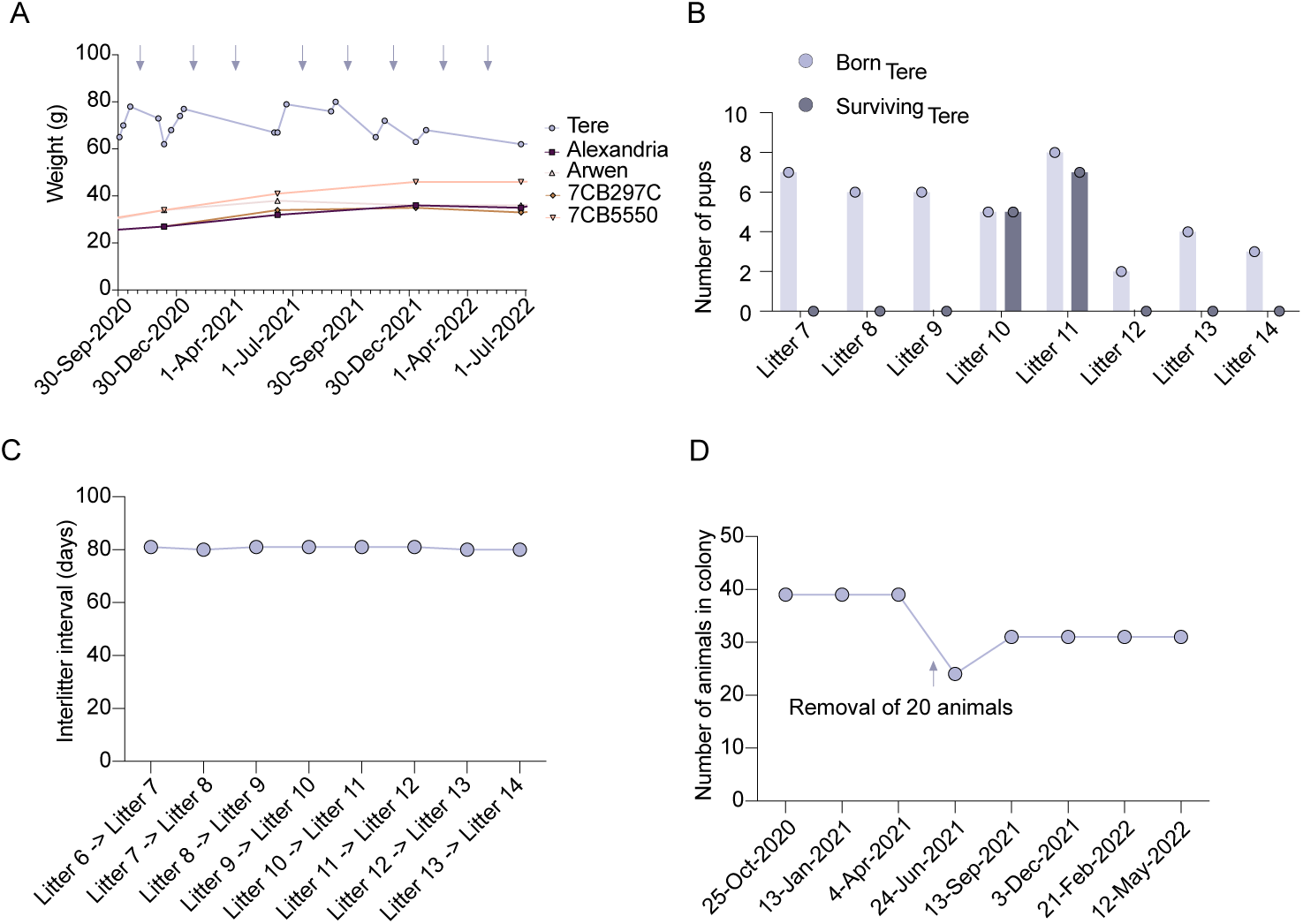
Environmental stressors impair the queen’s reproduction. (**A**) Body weight of Queen Teré and four of her daughters born in Queen Teré’s December 13, 2019 litter. Blue arrows indicate Queen Teré’s litters. (**B**) The number of pups born and the number of surviving pups past one month of age for the indicated litters. (**C**) Interlitter interval days between the indicated litters for the colony. (**D**) The number of animals in each colony one month following the noted litter dates. Data shown in panels **A**-**D** are also shown in **Supplemental** Figure 1A.

Finally, during these periods of reproductive instability no subordinate females exhibited signs of pregnancy or produced litters. Taken together, the temporal association between alleviated colony density and the resumption of viable births suggests that increased colony density can function as a stressor that compromises postnatal litter viability, but not disrupt the queen’s physiological capacity to conceive. Moreover, these data demonstrate that the decline in pup survival did not lift reproductive suppression among subordinates, consistent with patterns in other rodents in which elevated density inhibits, rather than promote reproduction.

### Reproductive disruption following relocation to a new facility

Relocation to new facilities is a well-established physiological and behavioral stressor in laboratory rodents. Relatively minor alterations in housing conditions such as changes in ambient room characteristics, novel olfactory profiles, or modifications of routine husbandry can significantly disrupt reproductive function (*16*). In mice and rats, interfacility transfers are associated with decreased fertility, early pregnancy loss, impaired maternal behavior and prolonged reproductive pause (*16*). Whether comparable environmental perturbations influence reproduction in captive naked mole rats has not been systematically examined. We therefore tested the hypothesis that transferring the Amigos colony would impair Queen Teré’s fertility, which could reveal forms of reproductive plasticity.

On May 23, 2022, we relocated the entire Amigos colony, along with their original housing, from the EBS vivarium to the SAF rodent facility at our institution. Environmental parameters including temperature, humidity, light/dark cycle and husbandry routine were matched between facilities. Nevertheless, following transfer, Queen Teré’s reproductive output ceased (**Figure 1**, **Table 1**, **Figure 4A** and **Supplemental Figure 1A, C-D**). Over the subsequent year, Teré maintained a stable body weight and serial clinical examinations revealed no morphological evidence of pregnancy (**Figure 4A** and **Supplemental Figure 1A**). Her next parturition did not occur until April 9, 2023 when she delivered a single pup that did not survive (**Figure 1**, **Table 1**, **Figure 4B**). Thus, despite controlled environmental equivalence between vivaria, the relocation was associated with prolonged reproductive arrest suggesting that movement to a new facility can compromise queen reproductive success by preventing normal litter production.

**Figure 4.**
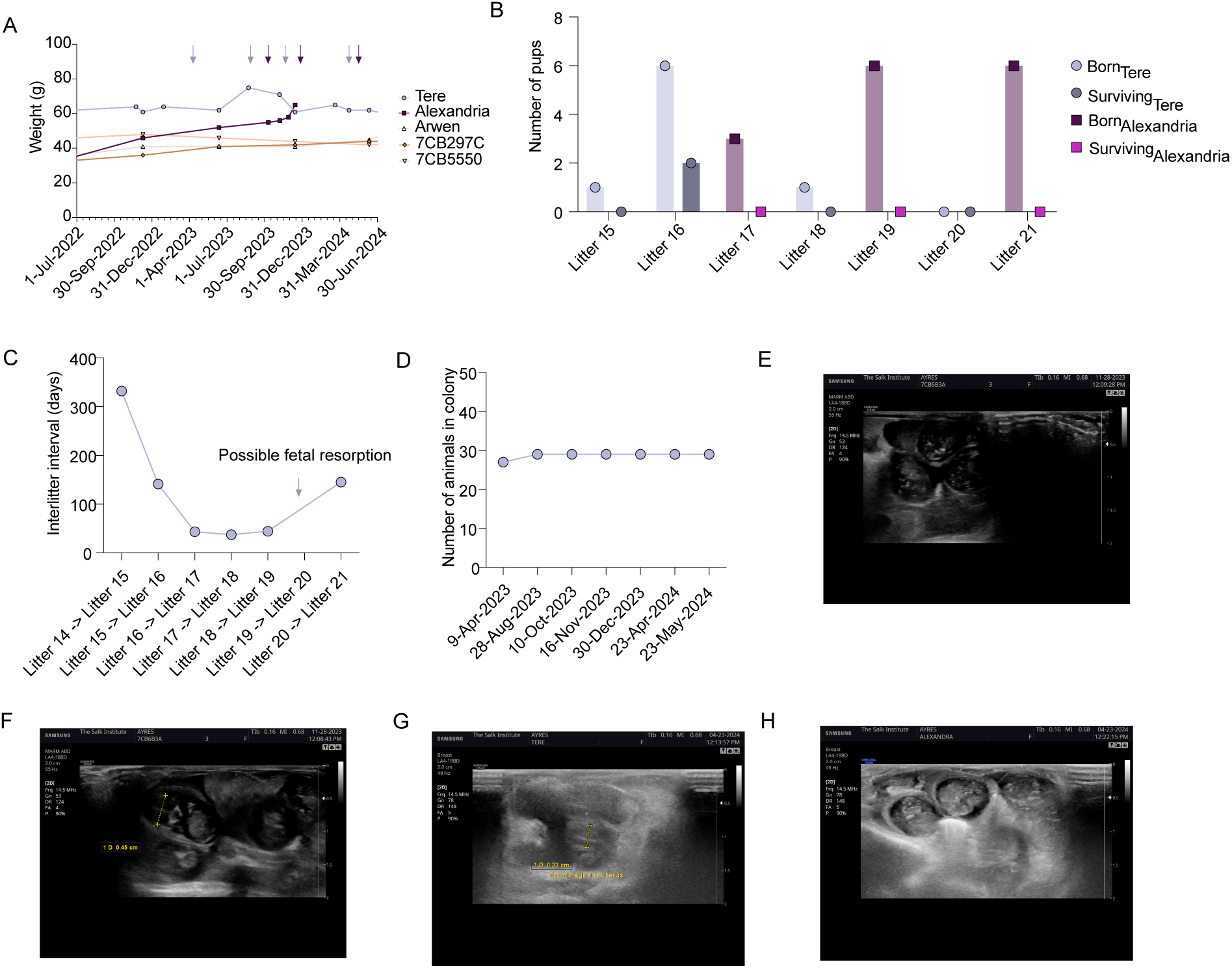
Emergence of a second reproductive female. (**A**) Body weight of Queen Teré and four of her daughters born in Queen Teré’s December 13, 2019 litter including Alexandria. Blue arrows indicate Queen Teré’s litters. Purple arrows indicate Alexandria’s litters. (**B**) The number of pups born and the number of surviving pups past one month of age for the indicated litters. Litter numbers indicate the litter for the colony. (**C**) Interlitter interval days between the indicated litters for the colony. (**D**) The number of animals in the colony one month following the noted litter dates. (**E-F**) Ultrasound imaging of Alexandria preceding her second litter (litter 19 for the Amigos colony). (**G**) Ultrasound imaging of Queen Teré in April 2024 indicating possible fetal resorption. (**H**) Ultrasound imaging of Alexandria in April 2024 preceding her third litter (litter 21 for the Amigos colony) confirming pregnancy. Data shown in panels **A**-**D** are also shown in **Supplemental** Figure 1A.

### Emergence of a second reproductive female

Following Queen Teré’s unsuccessful single-pup litter on April 9, 2023, she subsequently produced her 16^th^ litter on August 28, 2023, consisting of six pups of which two survived, with an interlitter interval of 141 days (**Figure 1**, **Table 1**, **Figure 4A-D** and **Supplemental Figure 1A-D**). 43 days later, on October 10, 2023, an additional litter of three pups was born, none of which survived (**Figure 1**, **Table 1**, **Figure 4B-C** and **Supplemental Figure 1A-B, D**). Because this interbirth interval was shorter than the 70-90 day gestation period of naked mole rats (*17*), we hypothesized that reproductive suppression may have been alleviated and a second reproductive female had begun cycling and contributing offspring. Around this time, we noted that one of Teré’s daughters from the December 13, 2019 litter exhibited morphological features consistent with reproduction including increased body length, more prominent abdominal contour and prominent nipples. Longitudinal body-weight trajectories confirmed that this animal displayed a distinct weight spike relative to her littermates, a signature that in our colony reliably precedes pregnancy (**Figure 4A** and **Supplemental Figure 1A**). We designated this female as Alexandria.

On November 16, 2023, Queen Teré produced another litter consisting of a single pup that did not survive (**Figure 1**, **Table 1**, **Figure 4A-B** and **Supplemental Figure 1A-B**). Concurrently, Alexandria exhibited a second discrete weight increase, indicating pregnancy, which we confirmed by ultrasound (**Figure 4A**, **E-F** and **Supplemental Figure 1A**). Alexandria subsequently delivered a litter on December 30, 2023 consisting of 6 pups with no survivors (**Figure 1**, **Table 1**, **Figure 4B** and **Supplemental Figure 1B**). During Spring 2024, Teré again displayed weight gain indicating possible pregnancy, however we did not detect any fetuses in Queen Teré during ultrasound examination (**Figure 4A, G** and **Supplemental Figure 1A**). This indicates that Queen Teré was never pregnant or that she was pregnant but underwent fetal resorption prior to the ultrasound examination. At this time, we also performed an ultrasound on Alexandria and confirmed that she was pregnant again (**Figure 4H**). On May 23, 2024, Alexandria gave birth to six pups, none of which survived (**Figure 1**, **Table 1**, **Figure 4B** and **Supplemental Figure 1B**). Within the week following this litter, Alexandria exhibited a rapid decline in activity level and overall condition. We performed humane euthanasia on May 29, 2024 and identified a uterine torsion as the proximate cause of decline from our necropsy analysis. Importantly, this plural reproductive state occurred without any observed aggression, dominance challenge or social instability (**Table 2**). Taken together, our data demonstrate that following a prolonged period of reproductive quiescence and multiple unsuccessful or minimal pup-litters from the established queen, a second reproductive female emerged and maintained pregnancies that were asynchronous yet partially overlapping with those of Queen Teré. Although reproductive suppression was clearly lifted in Alexandria, neither her litters nor the queen’s litters survived during this interval. Thus, the emergence of a second reproductive female did not restore net reproductive success to the colony within this timeframe.

### Peaceful queen succession

Following Alexandria’s euthanasia, Queen Teré produced one additional litter on September 9, 2024 yielding an interlitter interval for the colony of 109 days (**Figure 1**, **Table 1**, **Figure 5A-D** and **Supplemental Figure 1A-D**). This litter consisted of a single pup that did not survive, continuing the pattern of reduced reproductive success we observed throughout the preceding year (**Figure 1**, **Table 1**, **Figure 5B-C** and **Supplemental Figure 1B-C**). Shortly after this litter, another subordinate female, one of Teré’s daughters from the same litter as Alexandria (December 13, 2019 litter), began to display phenotypic changes consistent with reproductive activation including pronounced body weight increase and morphological changes (**Figure 5A** and **Supplemental Figure 1A**). Our ultrasound examination confirmed the presence of fetuses in this subordinate female that we named Arwen (**Figure 5E**). We observed no fetuses from our ultrasound imaging of Teré at this time.

**Figure 5.**
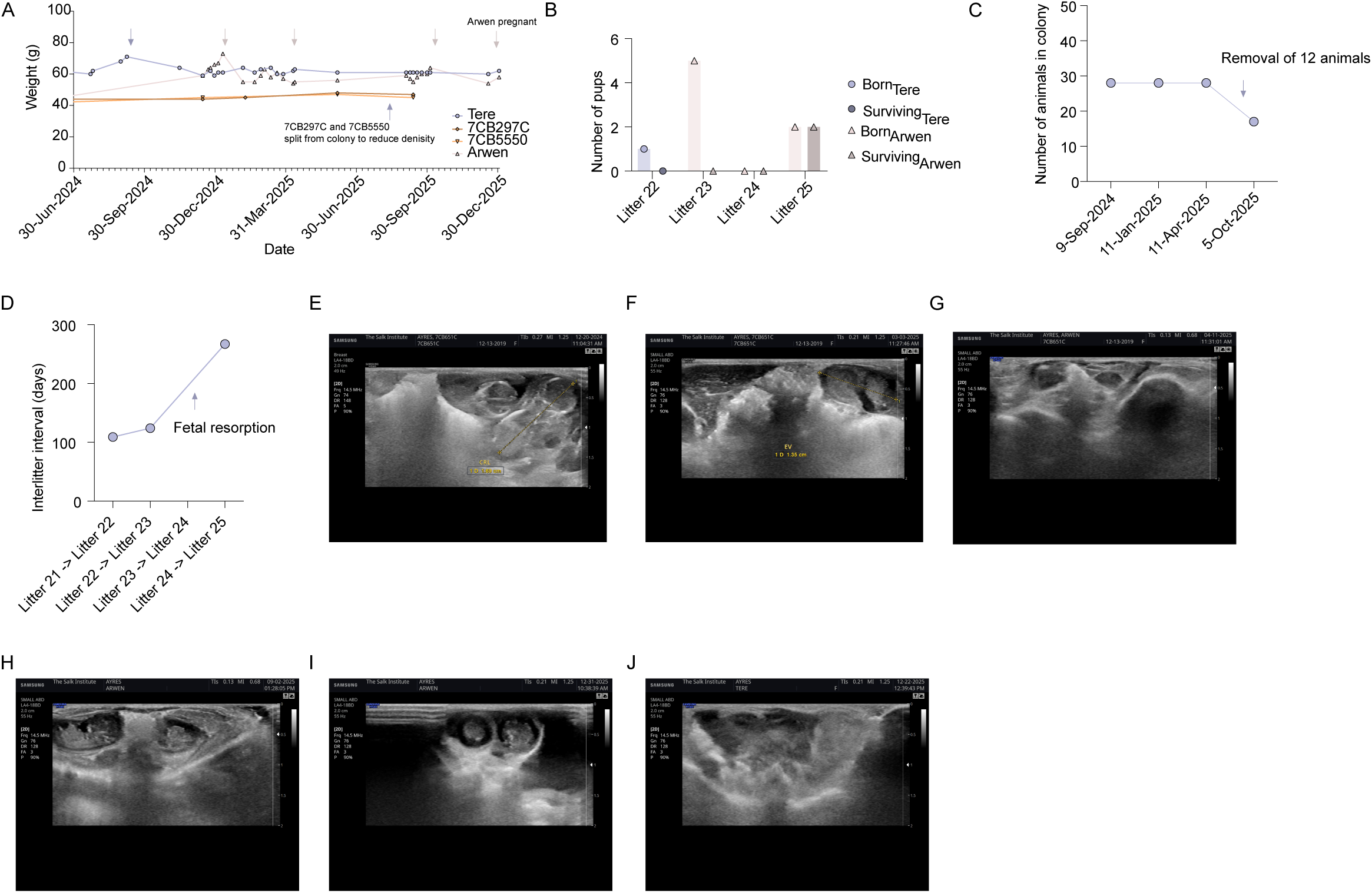
Peaceful succession of Queen Arwen. (**A**) Body weight of Queen Teré and three of her daughters born in Queen Teré’s December 13, 2019 litter including Arwen. Blue arrows indicate Queen Teré’s litters. Pink arrows indicate Arwen’s litters. (**B**) The number of pups born and the number of surviving pups past one month of age for the indicated litters. Litter numbers indicate the litter for the colony. (**C**) The number of animals in the colony one month following the noted litter dates. (**D**) Interlitter interval days between the indicated litters for the colony. (**E**) Ultrasound of Arwen’s first litter in December 2024. (**F**) Ultrasound of Arwen’s second litter in March 2025. (**G**) Ultrasound of Arwen’s second litter in April 2025 indicating fetal resorption. (**H**) Ultrasound of Arwen’s third litter in September 2025. (**I**) Ultrasound of Arwen’s fourth litter in December 2025. (**J**) Ultrasound confirming no pregnancy in Teré in December 2025. Data shown in panels **A**-**D** are also shown in **Supplemental** Figure 1A.

On January 11, 2025, Arwen delivered a litter of five pups, none of which survived beyond the early postnatal period (**Figure 1**, **Table 1**, **Figure 5B** and **Supplemental Figure 1A**). Arwen again exhibited clinical indicators of pregnancy in early Spring 2025 (**Figure 5A** and **Supplemental Figure 1B**) and an ultrasound performed in in March 2025 confirmed pregnancy (**Figure 5F**). Subsequently Arwen’s weight declined prompting us to perform another ultrasound on April 11, 2025 in which we no longer detected fetuses, suggesting fetal resorption occurred (**Figure 5G**). We hypothesized that the persistent neonatal mortality across multiple reproductive females might reflect density related limitations in postnatal care or worker provisioning. We therefore, removed 12 animals on September 11, 2025 to reduce the social group size while Arwen was pregnant with her third litter (litter 25 for the colony) (**Figure 5C, H** and **Supplemental Table 1**). Following this intervention, Arwen gave birth on October 5, 2025 to a litter of two pups, both of which survived, representing the first successful pup survival in the colony since 2023 (**Figure 1**, **Table 1**, **Figure 5B** and **Supplemental Figure 1B**). In December 2025, Arwen again displayed signs of pregnancy and ultrasound confirmed pregnancy with her fourth litter (**Figure 1**, **Table 1**, **Figure 5A**, **I** and **Supplemental Figure 1A**).

During this entire period of Arwen’s reproductive activity, Queen Teré showed no evidence of continuing fertility. Her body weight remained stable and ultrasounds failed to detect pregnancy at any point (**Figure 5A**, **J** and **Supplemental Figure 1A**). Aside from a single incident on February 6, 2025 in which one animal was found with a superficial bite wound and dried blood around the face, an injury that resolved without recurrence, no aggression or dominance related conflict was observed (**Table 2**). Instead, Queen Teré was reported to exhibit “guarding” behavior of Arwen and her litter. Importantly, no other signs of social instability, behavioral escalation or colony wide distress were documented. Taken together, these observations indicate that following the decline of Queen Teré’s reproductive capacity and the loss of the intermediary breeder Alexandria, Arwen successfully assumed the reproductive role without eliciting aggression from the reigning queen or from other colony members.

## Discussion

Naked mole rat queen succession is traditionally framed as a largely aggressive process. Here, we provide evidence that this canonical model does not fully capture the range of reproductive dynamics that are possible within naked mole rat colonies. By long-term longitudinal monitoring of a captive colony coupled with external stressors, we demonstrate that impairment of the queen’s reproductive success without a destabilization of the colony social structure can give rise to an alternative reproductive trajectory, involving peaceful plural breeding and queen succession. Our findings reveal a previously underappreciated plasticity in the mechanisms governing reproductive hierarchy, demonstrating that shifts in reproductive status of subordinates can occur in response to fertility-linked changes in the queen with colony social stability in addition to overt aggression and behavioral dominance.

The aggressive and rigid reproductive strategy of the naked mole rat is proposed to confer several fitness advantages under their harsh, but largely stable, natural environmental conditions (*18–20*). Restricting reproduction to a single queen minimizes reproductive conflict, reduces the risk of infanticide and ensures colony resources are concentrated on supporting one large litter rather than dispersed across multiple broods (*5, 21, 22*) By allocating their energy towards important tasks for the colony level such as tunnel excavations, foraging, defense and alloparental care, subordinates gain fitness benefits by supporting the queen’s offspring due to their high genetic relatedness (*23–26*). However, aggressive strategies can theoretically incur costs, especially when the queen’s reproductive output is disrupted. Indeed, in the banded mongoose, there were substantial health costs to pups born to dominate females that evicted subordinate females to trigger subordinate abortions (*27*). Peaceful plural breeding could buffer against a pause or failure of any single reproductive female and maintain colony cohesion, preserve potential breeders and lead to faster reproduction and colony growth. In agreement with some anecdotal reports (*28–32*), we found that in response to an external stressor that disrupted reproductive output of the queen, there was a period of peaceful plural breeding that ultimately led to the succession of a new reproductively successful queen. While we currently do not know the mechanistic rationale to explain when a colony follows the traditional aggressive reproductive trajectory versus the less common peaceful trajectory to queen succession, it is possible that under some conditions, the peaceful trajectory is favored because aggression-based enforcement may be insufficient or unnecessary, and when the cost of a “war” may be too high.

There are several models to explain how peaceful queen succession can occur. First, the reigning queen can halt reproduction and eventually a new female assumes queen status. Second, the reigning queen can halt reproduction, followed by plural breeding of subordinate females until one assumes the throne. Third, there can be a period of plural breeding involving the reigning queen and a subordinate female, with a subordinate female eventually assuming queen status. Consistent with the third model, we found that following a period of reproductive instability with social stability maintained, there was a transient successive phase of plural breeding involving Queen Teré and her daughter Alexandria. While plural breeding is uncommon in naked mole rats, anecdotal and limited empirical reports indicate that it can occur (*28–32*). Consistent with our study, these previous cases often involved closely related females, typically within the first few years following colony formation, was short lived and associated with small and asynchronous litters (*28–32*). In our study, plural breeding was characterized by poor pup survival and ended with the eventual nonviolent succession of Queen Arwen after we had to humanely euthanize Alexandria due to a uterine torsion. This is in contrast to previous reports in which plural breeding ended through escalated aggression or an unexplained disappearance of one breeder (*32*). Our data demonstrate that it is possible for previously reigning queens to peacefully assume nonreproductive roles in the colony.

Colony density is a well-established external stressor that can disrupt multiple aspects of reproductive physiology in rodent species. In mice and rats, overcrowding can diminish reproductive success via multiple mechanisms including increases in glucocorticoid levels, suppression of the estrous cycle and impairment of maternal behavior (*12–15*). These density dependent reproductive constraints can be observed in both wild and laboratory colonies, and can be exacerbated under captive settings (*33*). By contrast, in naked mole rats, high colony density is proposed to be foundational for their reproductive structure with the queen’s reproductive success being enabled by a large, high-density colony of non-breeding subordinates (*21*). In the current study, we found that increased density of a captive colony did not impair the queen’s ability to become pregnant and give birth to live litters at regular interlitter intervals. However, we did find that high colony density was associated with reduced pup survival and that alleviation of colony density led to increased pup survival. This is consistent with a previous report showing that pup survival decreased during the first ten days post birth in captive colonies with high density (*21*). Previous studies have reported that increased colony density is associated with heightened aggression and fighting in naked mole rat colonies, which is proposed to be the result of the queen’s need to reinforce her dominance and maintain the colony’s social hierarchy. (*5, 34*). In our study, we did not find increased colony density nor the associated impairment of pup survival to trigger aggression, fighting or any detectable disruption of social structure in our captive colony. While colony density is regulated by individuals dispersing to form new colonies in wild settings (*8*), in captive colonies, dispersal is not possible. Thus, our results likely reflect physical and behavioral constraints imposed by the artificial system rather than intrinsic reproduction limitations. Regardless, our system enabled us to demonstrate that impairment of queen reproductive success via reduced pup survival without triggering social instability was not sufficient to alleviate reproductive suppression of subordinate females in the colony.

Relocation to a new vivarium is a well-established stressor in laboratory rodents and is associated with reduced fertility, pregnancy loss, disrupted maternal behavior and temporary cessation of breeding (*35*). These effects are typically attributed to stress-induced elevations in corticosterone and other neuroendocrine changes triggered by alteration in sensory environment, handling and housing conditions (*36*). Despite identical environmental parameters and husbandry, relocation to a new facility disrupted the queen’s reproduction in the Amigos colony. Specifically, Queen Teré exhibited a complete pause in her reproductive output for almost one year following relocation. Importantly, during this time period, we observed no aggression, fighting or injuries consistent with fighting in our Amigos colony. While such relocations would not occur in the wild, this artificial perturbation allowed us to decouple the queen’s reproductive pause from social instability and test the effects on queen succession. Thus, laboratory environmental stressors can act as powerful experimental tools to uncover latent plasticity in the naked mole rat reproductive systems that may be masked in natural their habitat.

While our study was conducted on a single captive colony and relied on artificial environmental perturbations, our work provides an important conceptual advancement for our understanding of the naked mole rat social system. By uncoupling reproductive failure from social instability, we reveal an underappreciated mechanistic route for queen succession that does not involve aggression. Thus, our findings expand the framework for naked mole rat eusociality to include a “hidden” and non-violent flexibility for their reproductive reorganization. This plasticity may provide protection against periods of reproductive instability, enabling the animals to preserve their social cohesion while adapting to stressors. Future studies should focus on determining the natural conditions that trigger the peaceful route to queen succession in the wild.

## Acknowledgements

We thank members of the Ayres lab for helpful comments and suggestions. We thank Mat Leblanc and Sean Adams for helping to establish our naked mole rat colonies and protocols at Salk. We thank Dan McCloskey (The City University New York) for generously providing Queen Teré, Paquito and their first litter. We thank Trinka Adamson, Sarah Alaniz, Cassandra Brown, Sierra Garcia and Ashley Kliskey, for their input in colony split protocol, veterinary and husbandry care. This work was supported by the NOMIS Foundation and Howard Hughes Medical Institute (J.S.A.). Biorender was used for some figure panels.

## Materials and Methods

### Naked Mole Rat Colony – The “Amigos”

The original colony consisting of Queen Teré (DOB unknown), Paquito (DOB unknown) and their first litter (four females and one male born May 18, 2019) were generously provided by Dan McCloskey (CUNY). The colony arrived at the Salk Institute on July 17, 2019. All animals used in this study originated from this family. All experiments were performed in our AAALAC-certified vivarium, with approval from The Salk Institute Animal Care and Use committee (IACUC).

### Housing System

Upon initial arrival to the Salk Institute, the original 6 animals were placed briefly in a plastic Rubbermaid bin (23”L x 18”W x 15”H) with bedding and tubes placed below the bedding to serve as tunnels. They were then transferred to a 5 chamber plexiglass housing system shortly after (**Supplemental Figure 2A-B**). Briefly, the housing system consisted of five rectangular plexiglass chambers interconnected by cylindrical tubes mimicking burrow tunnels. At the end of each of these tunnels are sliding metal doors that can be used to block off or open the entrances. The top of each chamber consists of twelve holes around the circumference for circulation. By March 2020, the Amigos colony was placed in a new 10 chamber plexiglass housing but were only given access to five chambers. Two new chambers were opened in August 2020, giving them access to a total of 7 chambers. By the end of December 2020, the Amigos colony was utilizing the 10-chamber housing system, in which they are currently residing (**Supplemental Figure 2C**).

### Husbandry

All naked mole rats were housed in an animal facility room maintained at temperatures of 27 – 30°C, humidity between 50-100%, and a 17:7h dark-light cycle. If humidity levels were below 50%, deionized (DI) water was sprayed into the chambers. Light was provided 7h daily to allow for husbandry tasks, weighing, pregnancy examination, general observation and colony splits. Bedding was comprised of Pure-o’Cel and was replaced daily as needed. Enrichment included autoclaved cardboard pieces placed in the tunnels and throughout the housing for burrowing and chewing behaviors. Strips of paper towels dampened with DI water were provided daily for additional enrichment and maintenance of appropriate humidity levels within the system. Individual chambers were on a 6-week schedule to be removed and cleaned with DI water. Spot cleaning occurred daily as needed, especially for the bathroom chambers. Diet included raw yam and a daily rotation of supplemental corn, jicama, baby cereal, baby carrots, bell peppers, apple, and cabbage. For colonies of 10 or more animals, 9g yam and 5g of rotational items are provided per adult. For colonies with juveniles less than 1 year of age, 4.5g yam and 2.5g of rotational items are provided for each juvenile. In accordance with standard colony management, naked mole rats were not supplemented with water as dietary water content is sufficient to maintain appropriate hydration. In addition to investigator assessments, the veterinary staff checked all animals daily for general health and reproductive status.

### Facility Movement

The Amigos colony was housed in the Salk Institute’s EBS facility from June 2019 to May 2022. The colony was then moved to the SAF facility on May 23, 2022. All conditions in terms of lighting cycle, temperature and humidity were the same as described in the husbandry section above. They were maintained in their 10 chamber housing system. On October 26, 2023, the colony was moved back to the EBS facility.

### Weighing Protocol

The individual being weighed was temporarily removed from the colony and gently placed in a cleaned stainless steel bowl and placed on a weigh scale. After weight was noted, the animal was gently wiped with a paper towel from the colony housing prior to placement back into the colony to increase odor familiarity and prevent colony rejection of the individual. All individuals were weighed every six months unless otherwise indicated for health or pregnancy monitoring.

### Pregnancy Ultrasound Examination

Individual pregnancy examinations were performed with a Samsung HS60 ultrasound and a LA4-18BD linear transducer with a frequency range of 4 – 18 MHz. 2D ultrasonograms were obtained and measurements of embryonic vesicles and fetuses ranging from embryonic vesicle diameter, biparietal diameter, and crown – to – rump length were determined to estimate gestational age and parturition date as previously described (*17*). Females were temporarily taken out of the colony and gently held in dorsal recumbency by a veterinary technician. Approximately 5-10mL of warmed Aquasonic 100 Ultrasound Transmission Gel was applied to the abdomen. All ultrasound examinations were performed by a veterinarian within a period of 10 minutes to minimize restraint and stress of the animal. Each animal was then weighed and placed back in the colony as described in the Naked Mole-Rat Weighing Protocol.

### Animal Care Practices After Parturition

When a new litter was born, the colony was placed on a “Do Not Disturb” protocol for 30 days to increase likelihood of pup survival. Husbandry staff continued to remove leftover food and provide new food on a daily basis. The nesting chamber was not disturbed during this period; in addition, chamber removal for cleaning was paused for 30 days. Spot cleaning of chambers not including the nesting chamber was done on an as needed basis. Bedding was replaced as needed.

### Colony Split Protocol

Animals to be removed were placed in a new housing system. The new housing system was spot cleaned only with mild detergent and DI water to minimize stress in the separated individuals. No material from the original colony was placed into the new colony so that a new colony odor could be established. All individuals were weighed for baseline measurements as part of new colony healthy records. Food including base raw yam diet and supplements were recalculated based on the number of individuals in each colony.

## Supplemental Figure Legends

**Supplemental Figure 1.**
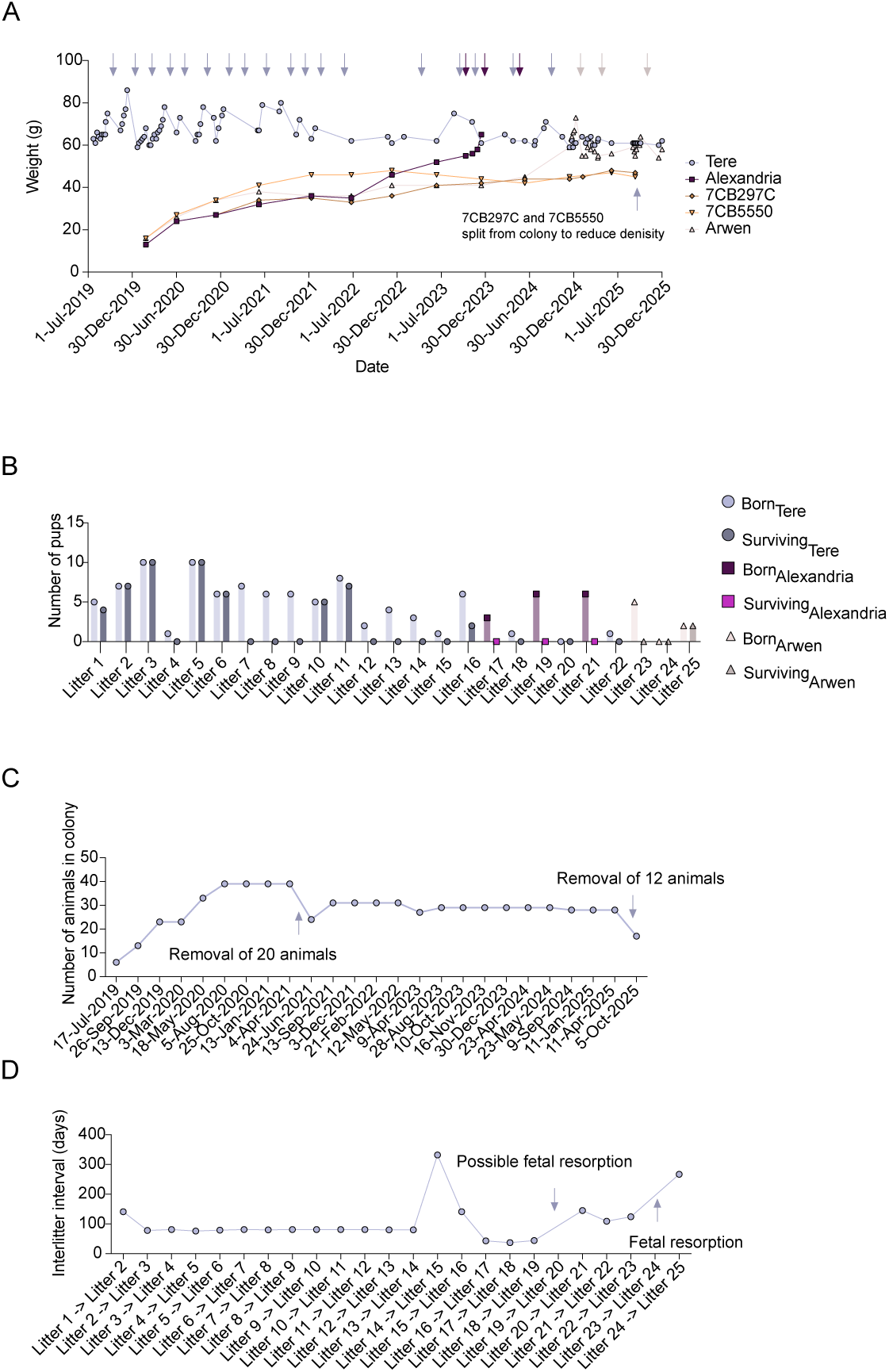
Summary of study data. (**A**) Body weight of Queen Teré and four of her daughters born in Queen Teré’s December 13, 2019 litter, including Alexandria and Arwen. Blue arrows indicate Queen Teré’s litters. Purple arrows indicate Alexandria’s litters and pink arrows indicate Arwen’s litters. (**B**) The number of pups born and the number of surviving pups past one month of age for the indicated litters. (**C**) The number of animals in the colony one month post noted litter dates. (**D**) Interlitter interval days between the indicated litters for the colony. Data from each panel are also displayed in **Figures 2-5**.

**Supplemental Figure 2.**
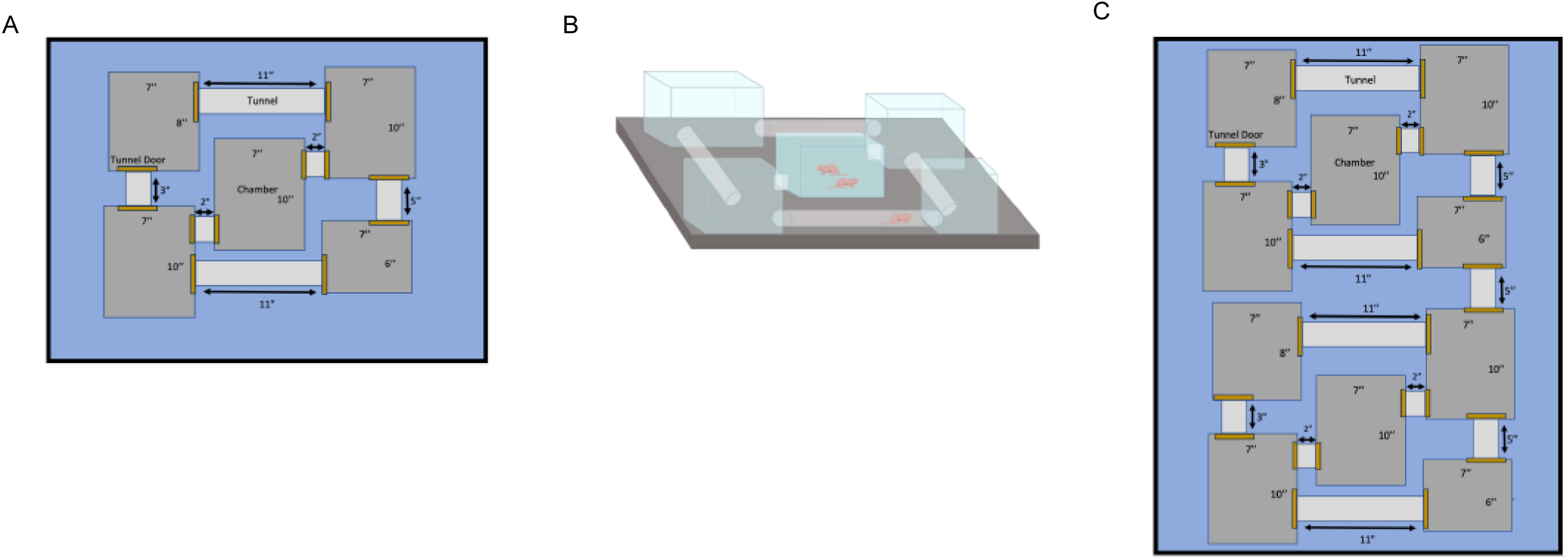
Housing used in this study. **(A)** 5 Chamber housing System. Chambers are in dark gray and range in dimensions of 7”W × 10”L, 7”W x 8”L, 7”W x 6”L. The height of all chambers is 7”. Tunnels are in light gray and range in lengths of 11”, 5”, 3”, and 2.” (**B**) 3D Model of 5 Chamber housing System. (**C**) 10 Chamber housing System. Chambers are in dark gray and range in dimensions of 7”W × 10”L, 7”W x 8”L, 7”W x 6”L. The height of all chambers is 7”. Tunnels are in light gray and range in lengths of 11”, 5”, 3”, and 2.”

## Supplemental Tables

**Supplemental Table 1.** Summary of animals removed from the Amigos colony over the course of the study.

| Animal ID | Birthdate | Sex | Date removed | Reason for Removal |
| --- | --- | --- | --- | --- |
| 7AA94B0 | 9/26/19 | F | 6/21/21 | Reduce Amigo colony density |
| 7CB643C | 12/13/19 | M | 6/21/21 | Reduce Amigo colony density |
| 7CB659D | 12/13/19 | M | 6/21/21 | Reduce Amigo colony density |
| 7CB6D80 | 12/13/19 | F | 6/21/21 | Reduce Amigo colony density |
| 7CB61CB | 12/13/19 | M | 6/21/21 | Reduce Amigo colony density |
| 7CB273C | 12/13/19 | F | 6/21/21 | Reduce Amigo colony density |
| 7CB6941 | 5/18/20 | M | 6/21/21 | Reduce Amigo colony density |
| 7CB6694 | 5/18/20 | M | 6/21/21 | Reduce Amigo colony density |
| 7CB480F | 5/18/20 | M | 6/21/21 | Reduce Amigo colony density |
| 7AAA1E6 | 5/18/20 | F | 6/21/21 | Reduce Amigo colony density |
| 7AAA8CBB | 8/5/20 | F | 6/21/21 | Reduce Amigo colony density |
| 7AA82F0 | 8/5/20 | M | 6/21/21 | Reduce Amigo colony density |
| 7AAA458 | 8/5/20 | M | 6/21/21 | Reduce Amigo colony density |
| 7BF324C | 8/5/20 | F | 6/21/21 | Reduce Amigo colony density |
| 7CB64B6 | 12/13/19 | F | 6/21/21 | Reduce Amigo colony density |
| 7CB4467 | 5/18/20 | F | 6/21/21 | Reduce Amigo colony density |
| 7CB616F | 5/18/20 | F | 6/21/21 | Reduce Amigo colony density |
| 7AA8F6A | 7/18/19 | F | 6/21/21 | Reduce Amigo colony density |
| 7AAABAC | 9/26/19 | M | 6/21/21 | Reduce Amigo colony density |
| 7BDC332 | 9/26/19 | M | 6/21/21 | Reduce Amigo colony density |
| Garbanzo | 9/13/21 | F | 7/8/22 | Start a new family |
| 7BDC31E | 5/18/20 | F | 7/8/22 | Start a new family |
| Reine | 7/18/19 | F | 7/15/22 | Start a new family |
| 7BD93DA | 9/13/21 | M | 8/18/22 | Missing - suspected escape |
| Alexandria | 12/13/19 | F | 9/9/24 | Euthanized - uterine torsion |
| 7BDC323 | 7/18/19 | F | 6/30/25 | Found dead - no fight wounds, nothing remarkable noted in autopsy. Cecum distended, postmortem change? |
| 7AAA327 | 9/26/19 | F | 9/11/25 | Reduce Amigo colony density |
| Bruiser | 9/26/19 | F | 9/12/25 | Reduce Amigo colony density |
| 7CB297C | 12/13/19 | F | 9/13/25 | Reduce Amigo colony density |
| 7CB5550 | 12/13/19 | F | 9/14/25 | Reduce Amigo colony density |
| 7BF31A7 | 9/26/19 | M | 9/15/25 | Reduce Amigo colony density |
| 7AAB632 | 8/5/20 | M | 9/16/25 | Reduce Amigo colony density |
| Pogo | 6/24/21 | F | 9/17/25 | Reduce Amigo colony density |
| 7BDEAA0 | 6/24/21 | F | 9/18/25 | Reduce Amigo colony density |
| 7BDC047 | 6/24/21 | F | 9/19/25 | Reduce Amigo colony density |
| 7BDE7C2 | 6/24/21 | M | 9/20/25 | Reduce Amigo colony density |
| 7BF2BCC | 9/13/21 | M | 9/21/25 | Reduce Amigo colony density |
| 7BDECB6 | 9/13/21 | M | 9/22/25 | Reduce Amigo colony density |

## Notes

### Competing Interest Statement

The authors have declared no competing interest.

### Summary of Updates

Updates to Figure 3. Updates to the text and figure legends.

